# Phosphatidylserine Exposure after Vascular Injury-Platelet or Endothelial

**DOI:** 10.1101/115139

**Authors:** Ejaife O. Agbani, Jenna L Cash, Christopher M. Williams, Alastair W. Poole

## Abstract

**Background:** Platelets membranes are considered the paramount site for the assembly of tenase and prothrombinase complex and are key players in localising coagulation to wound sites. However, the endothelium is also known to express phosphatidylserine (PS) and support the binding of recombinant FVa/FXa even beyond the site of injury. It thus remains unclear, what cell type play the preeminent role in the cellular control of coagulation after vascular injury.

**Approach:** To address this question, we utilised a model of haemostasis (full thickness 1mm excisional skin wounds) as well as tissues after injury in laser and ferric chloride models of thrombosis. Damage to the endothelium was assessed by the combined methods of picrosirius red staining, immunofluorescence and electron microscopy. Using multiphoton microscopy, we then compared the spatial distribution of PS on platelets and the endothelium.

**Results:** Platelets and detectable PS significantly co-localised compared with similar analysis of endothelial cell and exposed PS on wounded carotids arteries which was not significant. Point injury by laser induced restricted damage of the endothelium which was associated with limited platelets recruitment. In consistence with platelets response after FeCl_3_ injury, platelets exposed most of the PS detected at the wound edge where skin vessels were transected in our haemostasis model (Correlation Coeff. 0.78 +/− 0.12 vs 0.35 +/− 0.23).

**Conclusions:** We surmised that data from the different models support a paradigm of graded haemostatic response to vascular injury, in which full platelets response is limited to wound sites exposing the sub-endothelial matrix.

## INTRODUCTION

Advances in intra-vital microscopy and the fluorescent labelling of coagulation factors have enabled the visualisation of the coagulation cascade at wound sites. However, data emerging from these studies appear to challenge the dogma that platelets play a pre-dominant role in the provision of the requisite procoagulant surface for the cellular control of coagulation at wound sites^1-3^. Previous studies have used bound annexin-V (AnxV) as marker for the site specific localisation of activated coagulation factors X and V, FXa/FVa at injury sites^1, 4^ In models of laser-induced vessel injury, FVa and FXa have been localised on vessel wall and in the wound vicinity, with only a fraction of these co-localised with platelets. It is therefore thought that only a small percentage of platelets within a growing thrombus expose PS and support coagulation^4^. Furthermore, other blood cells^2, 3^ and indeed the activated endothelium have been reported in recent intravital studies, to provide the procoagulant surface required for coagulation^1, 5^.

It is therefore unclear, what cell type play the preeminent role in the cellular control of coagulation. To address this question, we utilised models of haemostasis and thrombosis, namely full thickness excisional wounds as well as tissues after laser or ferric chloride injuries. We performed electron and multiphoton microscopy and compared the spatial distribution of phosphatidylserine on platelets with the intimal layer of the vessel wall. Here, we show that activated platelets played a predominant role in the provision of procoagulant membrane at sites of damaged endothelium. Platelets role was dependent on whether the endothelium was activated and or breached after injury, and to what extent. While endothelial cells in the vicinity of FeCl_2_ injuries become activated and exposed lower level of PS, this was not associated with additional platelet recruitment as previously reported in laser injury models^1, 5^. We conclude that data from the different models support a paradigm of graded haemostatic response to vascular injury, in which full platelets’ response is limited to wound sites exposing the sub-endothelial matrix.

## METHODS

### Intravital microscopy of thrombus formation in vivo: Ferric chloride and laser and injuries

Mice were bred and experimental procedures performed under UK Home Office licence PPL30/2908, held by AWP. Mice were anaesthetised with ketamine 100 mg/kg (Vetalar V, Pfizer) and 10 mg/kg xylazine (Rompun, Bayer). Platelets were labelled by intravenous administration of 100 mg/kg Dylight-488 conjugated anti-GPIbβ antibody, 10 min prior to induction of thrombosis. For ferric chloride injuries, right carotid arteries were exposed and 2×1 mm 12% ferric chloride-soaked filter paper was placed on the arterial adventitia for 3 min. For laser-induced vessel injuries, the left cremaster muscle was exteriorized and spread flat over an optically clear coverslip and continuously superfused. Thrombi were induced in arterioles with a diameter of 25-35 μm by a nitrogen ablation laser (MicroPoint; Photonic Instruments), which was introduced through the microscope objective. Timelapse microscopy of the injury site for 20 min was performed and images processed using ImageJ. Background fluorescence values measured upstream of the injury site were subtracted from thrombus-specific fluorescence and data expressed as integrated densities.

### Excisional Cutaneous Wounding

#### Multiphoton imaging

Multiphoton imaging was used to detect the fluorophores and collagen by second harmonic generation. Endothelial cells were labelled Alexa647-CD31 (cyan blue; excited @ 1180), platelets were double-labelled with CD49b and GPIb-Alexa488 (green, excited @ 870nm), phosphatidylserine (PS) exposing membranes were identified by Alexa568-Annexin-V (AnxV, red, excited@1030), and collagen fibres at wound site were revealed by second harmonic generation (SHG, Deep Blue). The multiphoton systems is a Leica SP8 tandem scanning system with Spectra Physics Deep See (670-1300nm plus 1040nm) laser for multiphoton excitation and additional lasers for single photon excitation (Argon, 561 and 633nm). The system is equipped with several external non-descanned detectors (2PMT+2 HyD) for multiphoton imaging, internal detectors for confocal laser scanning microscopy (1PMT+2HyD) and a transmitted light detector.

#### Data analysis

Data were analysed using GraphPad Prism 7 (San Diego, CA). Statistical significance was determined by the Friedman test, followed by Dunn’s multiple comparison test.

## RESULTS

Using bound annexin-V (AnxV) as a marker for the site specific localisation of FXa/FVa at injury site^1, 5^, we localised the distribution of phosphatidylserine (PS) exposing membranes by labelling with Alexa568-Annexin-V, full thickness 1mm excisional skin wounds or carotid arteries after ferric chloride injury. Data exemplified by Figure 1A, showed PS exposure co-localised mainly with platelets at injury sites in the ferric chloride model. Consistent with previous finding^1, 5^ we observed low level PS exposure without adherent platelets at the vicinity of platelet aggregates, and in areas of the endothelium distant from the thrombus core (related to Movie S1). Furthermore, we examined several carotid arteries after FeCl_2_ injury to determine the spatial distribution of endothelial cells, platelets and bound Annexin-V. Global co-localisation analysis of the injuries and surrounding sites indicated that platelets exposed the most PS. Platelets and detectable PS co-localisation was significant compared with similar analysis of endothelial cell and exposed PS on wounded vessels which was not significant. (Fig.1B; related to Movie S2). As previously reported^6^, vessel treatment with 12% ferric chloride induced endothelial layer injury and exposure of sub-endothelial collagen (Fig.1C). Conceivably, we observed thrombus formation mainly in the region of collagen exposure (Figure 1A-i,v-vi; related to Movie S1). In turn, collagen imaging by second harmonic generation was enhanced at these wound sites due likely to denuded/damaged endothelium which may have improved photon penetration (Figure 1A-i,v-vi; related to Movie S1).

**Figure 1:**
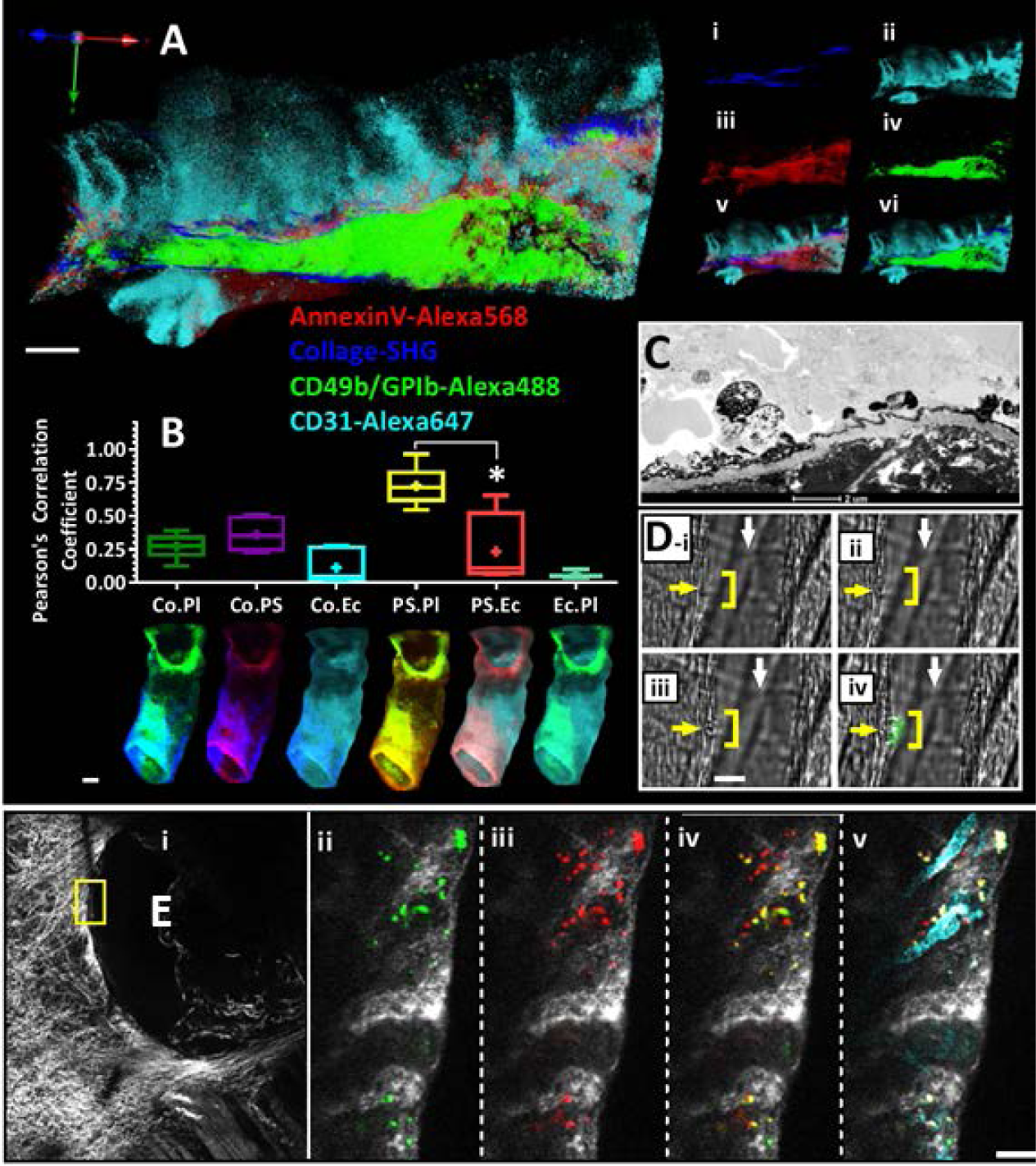
Phosphatidylserine (PS) exposure colocalised mainly with platelets at sites of endothelial damage in FeCl_3_ model of *in-vivo* thrombosis. **A:** Electron microscopy of a 30 min biopsy of mouse carotid artery following FeCl3 injury. **B:** 3-dimension reconstructions of multiphoton images of mouse carotid artery after FeCl_3_ injury. Endothelial cells were labelled Alexa647-CD31 (cyan blue), platelets were double-labelled with CD49b and GPIb-Alexa488 (green), PS exposing membranes were visualised by Alexa568-Annexin-V (AnxV, red), and collagen fibres at wound site were revealed by second harmonic generation (SHG, Deep Blue). Mini images in A (i-viii), show superimposed fluorescent images of background, collagen fibres, platelets and PS as indicated. **C:** Chart show summary of co-localisation analysis of fluorescent signal corresponding to PS, platelets (Pl) and endothelial cells (Ec) within carotid arteries. Carotid artery image corresponding to each analysis is shown on top row **D:** Cross-section of carotid artery taken after occlusive thrombus was process for fluorescent confocal microscopy. Platelets and PS were labelled as indicated in B and C; other blood cells are shown in grey. Data analysis was by Friedman test, followed by **E:** Platelets co-localised with phosphatidylserine and not endothelial cells at the wound edge in full thickness excisional wound model. Dunn’s multiple comparison test. *P* < 0.05 (*) was considered significant. Scale bar represent 2 pm (A), 100 pm (B, C, D). Data are representative of 1-5 mice.

Next, we supposed that platelets will localised mainly in the region of breached endothelium in our laser injury model. Indeed, point injury by laser induced limited damage of the endothelium as shown in Figure 1D-ii,iii; and this was associated with platelets recruitment to this site only (Figure 1D-iv). Although, activated endothelial cells exposing PS have been reported in the region around the point of laser injury^7^, in our study this did not cause platelets to adhere in these areas as shown in Fig.1D (related to Movie S3); this is also in consistence with recent reports^1, 5^. Finally, we examined the haemostatic response of the whole animal to full-thickness 1mm excisional wounding. In consistence with platelets’ response after FeCl_2_ injury, platelets exposed the most bound PS detected at the wound edge where small vessel were transected (Fig.1E; Correlation Coeff. 0.78±0.12 vs 0.35±0.23).

## DISCUSSION

Platelets′ membrane are considered the paramount site for the assembly of tenase and prothrombinase complex and a key player in localising coagulation to wound sites. However, the damaged or activated endothelium have now been shown to express PS and support the binding of recombinant FVa/FXa even beyond the site of injury^1, 5^. Notably, this observation was made in models of laser-induced vessel injury, where platelets assembly at the thrombus core corresponded to the point of laser injury. Also, report that platelet recruitment did not occur at regions showing FVa/FXa binding though distant from the wound site^1, 5^ raises questions about the spatial locale of thrombin generation especially since fibrin deposition co-localised with platelets in these models^4, 5^.

The interpretation of platelets role in the cellular control of coagulation is often model specific^5, 6, 8, 9^. A key factor to understanding platelets role after injury is whether the endothelium is activated and or breached after injury, and to what extent, in the particular model of haemostasis or thrombosis. Using FcRγ null mice (FcRγ^−/−^) lacking platelets surface collagen receptor glycoprotein VI (GPVI), Dubois et al., showed that while thrombus formation was significantly diminished in FcRγ^−/−^ compared with wild type controls after FeCl_3_ injury, the outcome was comparable after laser injuries^6^. Interestingly, collagen exposure was not detected at the site of laser injury by Dubois et al^6^; thus exposure of the sub-endothelial matrix may account for the difference in outcome^6^. Furthermore, a recent study by Chen^10^ and colleagues showed that vessel injury by FeCl_3_ resulted in an occlusive thrombus with profuse presence of procoagulant platelets, whereas very few procoagulant platelets were generated in the laser injury model and thrombus was non occlusive^10^.

In this study, we demonstrated a predominant role for platelets in the provision of procoagulant membrane after endothelial layer damage, where procoagulant agonists, of which collagen is the most abundant, is exposed. Logically, the mammalian system should balance the control of bleeding with the pathological consequences of thrombus formation. Therefore, activation of vessel intimal layer by heat shocks, pathogens or small injuries may induce endothelial PS exposure associated but with only minimal platelet response as exemplified by the laser injuries^5, 6^. Studies that demonstrate the primary dependence of stimulation laser injury model, on thrombin but not extra-vascular collagen for thrombus formation support this supposition^11, 12^. Fittingly, platelet mediated fibrin deposition may remain unchanged after pharmacological depletion but not obliteration of platelets^7, 9, 13^. Conversely, the damaged endothelium seen in FeCl_3_ injuries, exposes sub-endothelial collagen^6^ and evokes a GPVI mediated full haemostatic response in which platelets become activated and are recruited by an array of feedforward mechanisms^6, 14^. In conclusion, our data did not suggest a diminished role for platelets role in the regulation of coagulation after vessel injury. Instead, we surmised that data from the different haemostasis and thrombosis models support a paradigm of graded haemostatic response to vascular injury, in which full platelets’ response is limited to wound sites exposing the sub-endothelial matrix.

## Acknowledgements

We acknowledge the MRC and the Wolfson Foundation for funding the University of Bristol’s Bioimaging Facility. We thank Alan Leard, Katy Jepson and Judith Mantell of the Wolfson Bioimaging Facility for their assistance. We also thank Elizabeth Aitken for technical assistance. This work was supported by the British Heart Foundation (RG/10/006/28299) and the United Kingdom National Institute for Health Research (II-LB-0313-20003).

## Authorship contributions

EOA designed and performed experiments, analyzed data, contributed to discussion and co-wrote the manuscript. CMW and Jenna Davis-Cash performed *in vivo* experiments. IH supervised research, contributed to discussion and wrote the manuscript. AWP supervised research, contributed to discussion and co-wrote the manuscript.

